# Sex matters in Massive Parallel Sequencing: Evidence for biases in genetic parameter estimation and investigation of sex determination systems

**DOI:** 10.1101/096065

**Authors:** Laura Benestan, Jean-Sébastien Moore, Ben J. G. Sutherland, Jérémy Le Luyer, Halim Maaroufi, Clément Rougeux, Eric Normandeau, Nathan Rycroft, Jelle Atema, Les N. Harris, Ross F. Tallman, Spencer J. Greenwood, K. Fraser Clark, Louis Bernatchez

## Abstract

Using massively parallel sequencing data from two species with different life history traits -- American lobster (*Homarus americanus*) and Arctic Char (*Salvelinus alpinus*) -- we highlighted how an unbalanced sex ratio in the samples combined with a few sex-linked markers may lead to false interpretations of population structure and thus to potentially erroneous management recommendations. Multivariate analyses revealed two genetic clusters that separated males and females instead of showing the expected pattern of genetic differentiation among ecologically divergent (inshore vs. offshore in lobster) or geographically distant (east vs. west in Arctic Char) sampling locations. We created several subsamples artificially varying the sex ratio in the inshore/offshore and east/west groups, and then demonstrated that significant genetic differentiation could be observed despite panmixia for lobster, and that F_st_ values were overestimated for Arctic Char. This pattern was due to 12 and 94 sex-linked markers driving differentiation for lobster and Arctic Char, respectively. Removing sex-linked markers led to nonsignificant genetic structure (lobster) and a more accurate estimation of F_st_ (Arctic Char). We further characterized the putative functions of sex-linked markers. Given that only 9.6% of all marine/diadromous population genomic studies to date reported sex information, we urge researchers to collect and consider individual sex information. In summary, we argue that sex information is useful to (i) control sex ratio in sampling, (ii) overcome “sex-ratio bias” that can lead to spurious genetic differentiation signals and (iii) fill knowledge gaps regarding sex determining systems.

## Introduction

Recently, the revolution in massively parallel sequencing (MPS) technology has led to the production of many genome-wide datasets, whereby thousands of markers can be easily and inexpensively genotyped in hundreds of individuals for both model and nonmodel species (Davey *et al.* 2011; Andrews *et al.* 2016). Several MPS studies based on either RAD-sequencing or Genotype-By-Sequencing (GBS) techniques have demonstrated that these markers bring unprecedented insights on the causes and consequences of population structuring (reviewed in Narum *et al.* 2013). The strength of such methods comes from its supposedly random sampling of the entire genome (Davey *et al.* 2013). While the random distribution of markers achieved by these methods is advantageous in many regards, it has one over-looked result that could have consequences for inferences of population structure: some of the markers identified will be located on sex chromosomes, or in regions linked to sex, in species with genetic sex determination. Indeed, Wright (1931) pointed out this bias in genetic parameter estimations, particularly when sampling populations with varying sex ratios or in the presence of sex-biased dispersal. Despite the potential importance of these biases, few MPS studies have focused on the analysis of sex-linked markers (but see Gamble & Zarkower 2014; Kafkas *et al.* 2015; Brelsford *et al.* 2016; Larson *et al.* 2016) and to our knowledge, none have investigated the influence of sex-linked markers on inferences of population structure observed.

In addition to the importance of avoiding potential biases, detecting sex-linked markers in MPS datasets can also provide valuable information on sex determination (Pan *et al.* 2016). Sex is common to almost all living animals and often leads to the evolution of male and female dimorphism, both at the genetic and phenotypic level (Bell 1982). Diverse mechanisms acting at the scale of the genome, chromosomes or cells underlie the morphological, physiological and behavioral differences between males and females. Moreover, sex determination systems vary tremendously among and within taxa (Bachtrog *et al.* 2014), highlighting the challenges in determining the selective forces driving sex determination. In general, the diversity of sex determination systems reported in fish (particularly teleosts) and crustaceans is much more pronounced than that observed in mammals and birds (Bachtrog *et al.* 2014). Yet, the characterization of the genetic architecture of sex determination in these taxonomic groups has been limited to a few studies (Legrand *et al.* 1987). The access to new genomic approaches, which are increasingly being used in non-model marine and aquatic organisms (Kelley *et al.* 2016), offers new prospects to investigate the molecular basis of sex determination in this diverse group.

The identification of sex-linked markers can also provide a wealth of other useful information for management, conservation, and aquaculture (Pan *et al.* 2016). First, sex-linked markers can assist in the identification of the sex of an individual, particularly in cases with an absence of clear sexual dimorphism (*e.g.*, at young life history stages). In aquaculture practices, this can help farmers to maintain equal sex ratios of breeding adults and to implement efficient breeding programs (Martínez *et al.* 2014). Second, sex information is often important to include as a covariate in genetic models for finding loci linked to specific traits in order to reduce residual variation (Broman & Sen 2009). Third, knowing the sex of individuals may facilitate the demonstration of sex-biased dispersal, *i.e.*, when individuals of one sex are more prone to disperse (Prugnolle & De Meeûs 2002). Sex-biased dispersal is widely spread among vertebrates and can have important ecological and evolutionary consequences, but there is still little research on this topic in marine organisms, such as fishes and crustaceans, compared to mammals and birds (Mossman & Waser 1999).

Here, we present two empirical examples that illustrate how an unbalanced sex ratio combined with a few sex-linked markers can lead to false interpretations of population structure and to erroneous management recommendations, especially in species with high connectivity as frequently observed in marine and diadromous organisms. Our initial goal was to separately investigate population structure between two groups of American lobsters (*Homarus americanus*) occupying inshore and offshore habitats, and between Arctic Char (*Salvenius alpinus*) collected from two geographically separated regions (east and west) in the Canadian Arctic. In both cases, preliminary multivariate analyses mainly revealed two genetic clusters corresponding to male and female individuals instead of being related to inshore/offshore groups of lobsters or to east/west groups of Arctic Char. To further understand the clustering, we identified sex-linked markers driving the genetic differentiation between male and female in American lobster and Arctic Char. To demonstrate the potential impacts of sex-linked markers on the population genetic analysis, we tested for both species how different numbers of sexinked markers and ratios of samples from each sex can cause biased inferences of population structure. Finally, using the set of sex-linked markers identified, we found potential candidate genes or chromosomal regions linked to sex for American lobster and Arctic Char. We conclude with an exhaustive literature search demonstrating that very few studies performed on marine and diadromous species report sex information, and we argue, in light of our findings, that collecting this information can be critical to avoid biases, especially in high gene flow species.

## Methods

### Sampling and molecular techniques

American lobster: Commercial fishers collected 203 American lobsters (100 males and 103 females) from 13 sites including eight inshore sites and five offshore sites along the Atlantic coast of North America (Figure 1A; Table S1). The sex of all specimens was determined visually from obvious external morphological differences. Genomic DNA was then extracted using Qiagen Blood and Tissue kits. DNA quality was confirmed using visual inspection on 1% agarose gel followed by quantification with Quantit Picogreen dsDNA assay kits. RAD-sequencing libraries were prepared following the protocol from Benestan *et al.* (2015). Each individual was barcoded with a unique six-nucleotide sequence and 48 individuals were pooled per library. Real-time PCR was used to quantify the libraries. Single-end, 100 bp sequencing was performed on an Illumina HiSeq2000 platform at the Genome Québec Innovation Centre (McGill University, Montréal, Canada).

**Figure 1.**
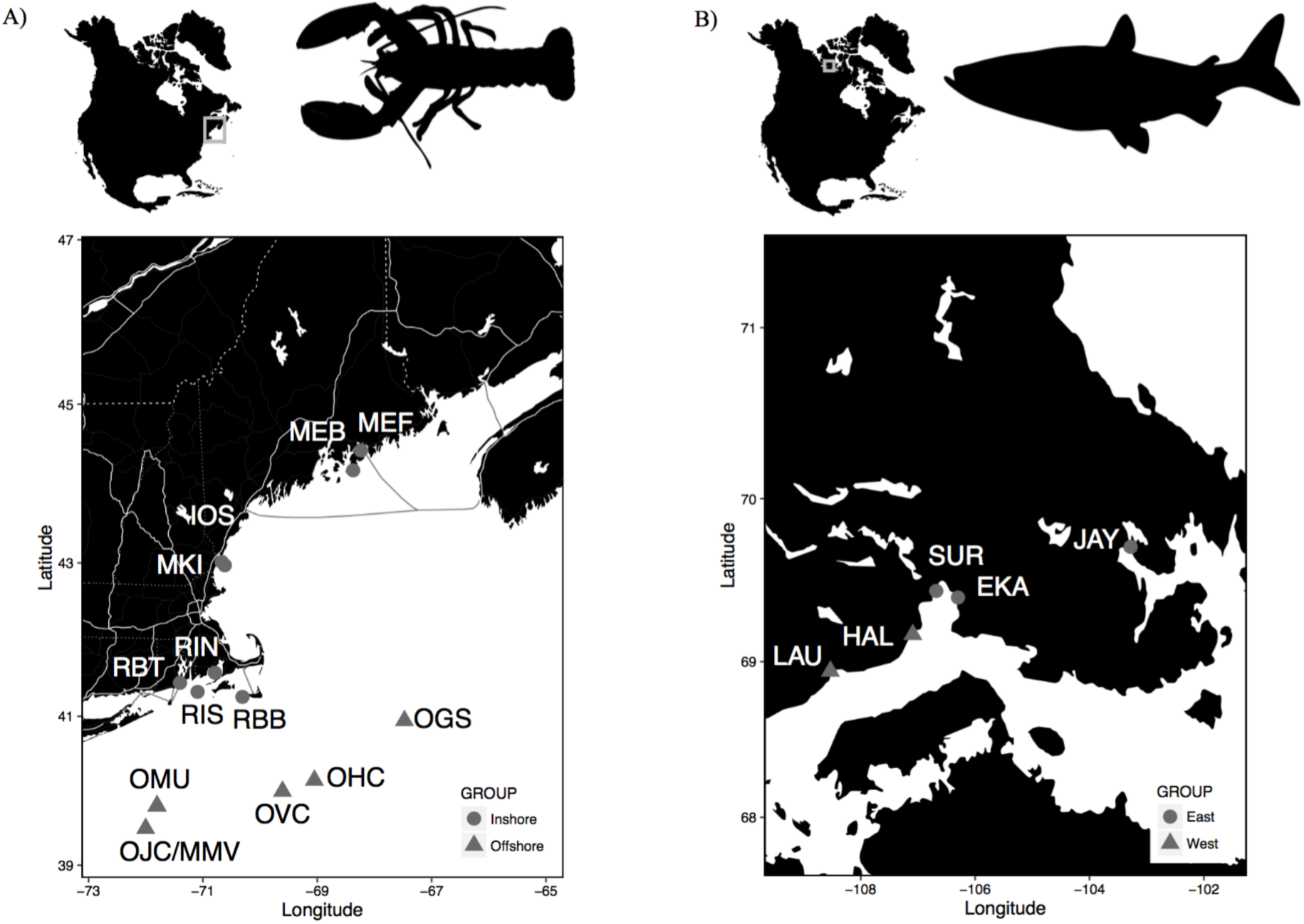
**Sampling Locations for American lobster (A) and Arctic Char (B).** (A) Inshore sampling locations are shown with a grey circle and offshore locations with a grey triangle. Inshore locations are Isle of Shoals (IOS; n=14), Blue Hill Bay (MEB; n=20), Frenchmans Bay (MEF; n=19), Kittery (MKI; n=20), Brown’s bank (RBB; n=17), Beavertail (RBT; n=16), Narragansett Bay (RIN; n=13), Rhode Island Sound Bay (RIS; n=7). Offshore locations are Georges Basin (OGS; n=10), Hydrographers Canyon (OHC; n=16), Jones Canyon (OJC; n=12), MacMaster Canyon (OMU; n=10) and Veatch Canyon (OVC; n=16). (B) Eastern sampling locations are shown with a grey circle and Western locations with a grey triangle. Eastern locations are Ekalluk (EKA; n = 58), Jayko (JAY; n = 58), Surrey (SUR; n = 30). Western locations are Halovik (HAL; n = 87) and Lauchlan (LAU; n = 57).

Arctic Char: Samples of 290 adult anadromous Arctic Char (142 males and 148 females) were collected from six rivers located on southern Victoria Island, Nunavut, Canada (Figure 1B; Table S2). Sex was determined visually by observation of the gonads for a subset (n = 174) and based on a genetic assay for another subset (n = 116), as described in Moore *et al.* (2016). In brief, the genetic sex was inferred based on the PCR assay described in Yano *et al.* (2013). Six individuals of known sex (three males and three females) were used as controls. Genomic DNA was extracted using a salt-extraction protocol modified from Aljanabi and Martinez (1997). DNA quality and quantity were checked on 1% agarose gels and using PicoGreen assays (Fluoroskan Ascent FL, Thermo Labsystems), respectively. Libraries were prepared based on a GBS protocol modified from Mascher *et al.* (2013) using *PstI* and *MspI* (details can be found in Perreault-Payette *et al.* in press). Specimens were individually barcoded with unique six-nucleotide sequences and pooled with 48 individuals per library. Libraries were each sequenced on two Ion Torrent Proton P1v2 chips.

### Bioinformatics and genotyping

Both the American lobster and Arctic Char libraries were de-multiplexed using *process_radtags* in STACKS (v.1.29 for American lobster and v.1.40 for Arctic Char) (Catchen *et al.* 2013). Raw sequencing data was checked in FASTQC (Andrews 2015). Reads were truncated to 80 bp for lobster and 70 bp for Arctic Char and adapter sequences were removed with CUTADAPT (Martin 2011).

American lobster: Loci were identified allowing a maximum of three nucleotide mismatches (M = 3), according to Ilut *et al.* (2014) and a minimum stack depth of three (m = 3), among reads with potentially variable sequences *(ustacks* module in stacks, with default parameters). Then, reads were clustered *de novo* to create a catalogue of putative RAD tags (*cstacks* module in STACKS, with default parameters). In the *populations* module of STACKS v.1.29 and following consecutive filtering steps, SNPs were retained when they were genotyped in at least 80% of the individuals and found in at least 9 of the 12 sampling sites. Potential paralogs were excluded by removing markers showing heterozygosity > 0.50 and 0.30 < F_IS_ < −0.30 within sites. Only SNPs with a global minor allele frequency > 0.02 were retained for the analysis. The resulting filtered VCF files were converted into the file formats necessary for the following analyses using PGDspider v.2.0.5.0 (Lischer & Excoffier 2012).

Arctic Char: SNPs were identified by first mapping the reads to the genome of the closely related Rainbow Trout *(Oncorhynchus mykiss;* Berthelot *et al.* 2014) using GSNAP v2016-06-09 with a minimum of 90% read coverage (-min-cov 90), tolerating 2 mismatches (-m 2) and setting an indel penalty to 2 (-i 2). A subsequent trimming step was conducted with SAMtools v1.2 (Li et al., 2009) to remove unmapped and multimapped reads using flags –F 1797 and –F 4, and a minimum mapping quality (MAPQ) of 1, respectively. The binary alignment files (bam) were then used as input for downstream analysis. Genotypes were obtained using STACKS v.1.40 integrated in a workflow developed in our laboratory (Benestan *et al.* 2016a). The catalog of loci was created allowing no mismatches among loci in *cstacks* (n=0) and a minimum stack depth of four (-m 4). SNPs were retained if at least 50% of the individuals were genotyped for the marker (-r 0.5) and the locus was present in at least four populations (-p 4). Potential paralogs were excluded by removing markers showing heterozygosity > 0.60 and F_IS_ - 0.40 F_IS_ > 0.40 within samples. Only SNPs with a global minor allele frequency > 0.01 were retained for the analysis.

### Discriminant Analysis of Principal Component (DAPC)

For American lobster, Discriminant Analysis of Principal Components (DAPC) was performed in the R package *adegenet* (Jombart *et al.* 2010). The optimal number of discriminant functions (n=60) to retain was evaluated according to the optimal α-score obtained from the data (Jombart *et al.* 2010). For Arctic Char, a Principal Component Analysis was performed in *adegenet.* As population differentiation was pronounced enough to be observed with the PCA, a DAPC was not conducted.

### Sex outlier loci detection

American lobster and Arctic Char: Outlier loci corresponding to the most divergent markers between sexes were identified with a level of differentiation between sexes exceeding random expectations using F_st_-based outlier analyses. Outlier SNPs were detected with BAYESCAN v. 2.1 (Foll & Gaggiotti 2008). BAYESCAN runs were implemented using permissive prior model (pr_odds) of 10, including a total of 10,000 iterations and a burn-in of 200,000 steps. For both species, these outlier analyses were conducted on the entire data set separated by sex.

### Sex ratio and sex-linked marker influence on index of genetic differentiation (Fst)

To determine the extent to which differing sex ratio influences the detected genetic structure, different proportions of male and female American lobsters or Arctic Char were subsampled from inshore or east and offshore or west, respectively, keeping a total of 50 individuals per group. This generated a gradient of six different sex ratio datasets, representing different sampling bias scenarios, from the most balanced (sex ratio = 25:25/25:25) to the most unbalanced sex ratio (sex ratio = 0:50/50:0).

Considering the three most unbalanced sex-ratio datasets (*i.e.*, 0:50/50:0, 5:45/45:5, 10:40/40:10), we removed sex-linked markers (*i.e.*, here outlier SNPs) according to their F_st_ values (in descending order) and we estimated F_st_ between offshore/inshore for the American lobster and east/west for the Arctic Char. We calculated F_st_ values using the function fst_WC84 in *assigner* R package (Gosselin *et al.* 2016).

### Marker annotation and genomic position

American lobster: There is no reference genome or high-density linkage map available for American lobster and so the approximate locations or associated linkage groups of the sex-linked SNPs could not be determined. Probable proximity between markers was determined by linkage disequilibrium (LD) analysis by calculating LD between pairs of SNPs using the *geno-r2* command available in VCFTOOLS (Danecek *et al.* 2011). The LD data frame obtained with VCFTOOLS was then transformed into an LD matrix to be analyzed using the *heatmap* command in the R environment (Team 2013). In order to determine what genes are associated with these sex-linked markers, the 12 candidate SNPs (outliers identified by BAYESCAN) were queried using BLAST against the transcriptome of the American lobster (F. Clark and S. Greenwood, University of Prince Edward Island, *personal communication;* see details in Benestan *et al.* 2016b). Six of the 12 candidate SNPs were distributed among six different contigs in the transcriptome data. The associated contigs were used as queries in a BLAST search against the SWISSPROT database (Bairoch & Apweiler 2000). A minimal *E*-value threshold of 1 × 10^−6^ and percent similarity of at least 70% were used. This yielded a set of two candidate SNPs associated with known genes. Gene ontology (GO) annotation terms were then associated to the candidate SNPs using SWISS-PROT accessions.

Arctic Char: There is no reference genome available yet for Arctic Char, but there is a high-density linkage map available for the closely related Brook Char (Sutherland *et al.* 2016). To obtain approximate positions of the sex-linked SNPs from Arctic Char, the MapComp method (Sutherland *et al.* 2016) was used to pair all of the Arctic Char markers with mapped Brook Char markers using the Atlantic Salmon genome (Lien *et al.* 2016; GenBank: GCA_000233375.4) as the intermediate reference genome. This method connects markers from two different linkage maps by mapping the markers to a reference genome, then pairing markers that map uniquely to the same place or close to each other in the reference genome. This was done as previously described (Sutherland *et al.* 2016), but with ten iterations to permit more than one anonymous marker pairing with a single mapped marker, as previously described (Narum *et al. in review)* but with a 1 Mbp maximum distance between the paired markers on a reference genome. This yielded approximate positions for determining the number and identity of linkage groups associated with sex in Arctic Char. To determine which genes are associated with these linked markers, the sex-linked markers were used in a BLAST query against the annotated Atlantic Salmon genome (Lien *et al.* 2016); NCBI Genome ICSASG_v2 reference Annotation Release 100).

### Literature search for marine and diadromous species population genomic studies

We performed an exhaustive literature search to document the proportion of population genomics studies that have reported sexing the species analyzed. More specifically, we conducted a literature search of population genomics studies on marine/diadromous species published in peer-reviewed journals from January 2010 to 15 November 2016 using the ISI Web of Knowledge bibliographic database (Thomson Reuters, http://thomsonreuters.com) using search keywords (i) “genomics” AND “marine” AND “SNP” yielded 22 hits, (ii) “population structure” AND “marine” AND “SNP” yielded 47 hits, (iii) “RAD-sequencing” AND “marine” yielded 39 hits and (iv) “population genomics” AND “marine” yielded 243 hits, (v) “population genomics” AND “anadromous” OR “catadromous” yielded 11 hits. From these hits, several criteria were used to determine which studies to include in our analyses. First, the paper needed to focus on a marine animal and use a set of more than 1,000 SNP markers. Second, the paper needed to refer to population genomics or related areas such as outlier identification because these are the target areas of research likely to be influenced by the sex ratio bias in sampling. After removing studies on non-marine or non-animal organisms, or those with too low density of markers, a total of 38 and 14 publications were retained for marine and diadromous species, respectively (listed in Table 1 and 2).

**Table 1.** Marine population genomics studies describing the name of the study, the organism studied, the method used to produce the genetic markers, the number of individuals sampled (N), the number of genetic markers (SNPs), the number of individuals sampled per location (N_POP_), the index of genetic differentiation observed among the location studied (F_st_).

| Study | Organism | Method | N | SNPs | Sex | $N_{POP}$ | $F_{st}$ |
| --- | --- | --- | --- | --- | --- | --- | --- |
| Araneda *et al.* 2016 | *Mytilus chilensis* | RAD-seq | 220 | 1,240 | No | 25-39 | 0.005 |
| Benestan *et al.* 2015 | *Homarus americanus* | RAD-seq | 586 | 10,156 | Yes | 30-36 | 0.0018 |
| Benestan *et al.* 2016b | *Homarus americanus* | RAD-seq | 562 | 13,688 | Yes | 30-36 | 0.0018 |
| Berg *et al.* 2015 | *Gadus morhua* | SNP-array | 194 | 8,809 | No | 8-48 | 0.0002-0.0709 |
| Berg *et al.* 2016 | *Gadus morhua* | SNP-array | 141 | 8,168 | No | 42-51 | 0.00123-0.0008 |
| Boehm *et al.* 2015 | *Hippocampus erectus* | RAD-seq | 23 | 11,708 | No | 5-9 | 0.0454-0.1012 |
| Bruneaux *et al.* 2013 | *Gasterosteus aculeatus* | RAD-seq | 288 | 6,834 | Yes | 48 | Unkn. |
| Cammen *et al.* 2015 | *Tursiops truncatus* | RAD-seq | 156 | 7,431 | No | 12-26 | Unkn. |
| Chu *et al.* 2014 | *Nucella lapillus* | RAD-seq | 30 | 4,000 | No | Unkn. | 0.0004-0.0474 |
| Corander *et al.* 2013 | *Clupea harengus* | RAD-seq | 2* | 4,756 | No | 6 | 0.005 |
| Ferchaud *et al.* 2014 | *Gasterosteus aculeatus* | RAD-seq | 60 | 33,993 | No | 20 | 0.056-0.111 |
| Ferchaud & Hansen 2016 | *Gasterosteus aculeatus* | RAD-seq | 177 | 28,888 | No | 20 | 0.002-0.458 |
| Galindo *et al.* 2010 | *Littorina saxatilis* | 454 seq | 30 | 2,454 | Yes | 15 | 0.03 |
| Gleason & Burton 2016 | *Chlorostoma funebris* | RAD-seq | 90 | 1,861 | No | 15 | 0.0042 |
| Guo *et al.* 2015 | *Gasterosteus aculeatus* | RAD-seq | 10* | 30,871 | No | 36 | 0.0282 |
| Hohenlohe *et al.* 2010 | *Gasterosteus aculeatus* | RAD-seq | 100 | 45,000 | No | 20 | 0.0020-0.1391 |
| Jackson *et al.* 2014 | *Epinephelus striatus* | RAD-seq | 620 | 4,234 | No | 14-32 | 0.002 |
| Lal *et al.* 2016 | *Pinctada margaritifera* | RAD-seq | 156 | 5,243 | No | 32-50 | 0.046 |
| Lamichhaney *et al.* 2012 | *Clupea harengus* | RNA-seq | 400 | 440,817 | No | 50 | Unkn. |
| Miller *et al.* 2016 | *Haliotis rubra* | GBS | 80 | 1,180 | No | 10 | 0.003 |
| Le Moan *et al.* 2016 | *Engraulis encrasicolus* | RAD-seq | 128 | 5,638 | No | 24-64 | Unkn. |
| Moura *et al.* 2014 | *Orcinus orca* | RAD-seq | 115 | 3,281 | No | 6-21 | 0.0346-0.334 |
| Nayfa & Zenger 2016 | *Pinctada maxima* | SNP-array | 85 | 1,130 | No | 25-33 | -0.0047-0.004 |
| Pecoraro *et al.* 2016 | *Thunnus albacares* | RAD-seq | 100 | 6,772 | No | 10 | 0.0273 |
| Picq *et al.* 2016 | *Hypoplectrus spp* | RAD-seq | 126 | 97,962 | No | 13-43 | 0.0042 |
| Poćwierz-Kotus *et al.* 2015 | *Gadus morhua* | SNP-array | 95 | 7,944 | No | 26-40 | 0.034 |
| Reitzel *et al.* 2013 | *Nematostella vectensis* | RAD-seq | 30 | 2,759 | No | 4-7 | 0.286-0.622 |
| Rodríguez-Ezpeleta *et al.* 2016 | *Scomber scombrus* | RAD-seq | 122 | 29,394 | No | 15-29 | 0.0157-0.039 |
| Sodeland *et al.* 2016 | *Gadus morhua* | SNP-array | 378 | 9,187 | No | 43-48 | 0.000-0.0189 |
| Stockwell *et al.* 2015 | *Scarus niger* | RAD-seq | 81 | 4,253 | No | 24-30 | 0.007 |
| Xu *et al.* 2016 | *Bathymodiolus platifrons* | RAD-seq | 28 | 9,307 | No | 10-18 | 0.0126 |
| Zhang *et al.* 2016 | *Larimichthys polyactis* | RAD-seq | 24 | 27,556 | No | 12 | < 0.001 |
| Bradbury *et al.* 2010 | *Gadus morhua* | EST seq | 300 | 1,641 | No | 15-26 | Unkn. |
| Jones *et al.* 2012 | *Gasterosteus aculeatus* | SNP-array | 121 | 1,159 | No | 4 o 6 | 0.031-0.383 |
| Therkildsen *et al.* 2013 | *Clupea harengus* | EST seq | 508 | 1,047 | No | 14-37 | 0.000-0.086 |
| Tepolt & Palumbi 2015 | *Carcinus maenas* | EST seq | 84 | 10 809 | No | 12 | 0.003-0.134 |
| Bay & Palumbi 2014 | *Acropora hyacinthus* | EST seq | 23 | 15,399 | No | 10-13 | Unkn. |
| De Wit & Palumbi 2013 | *Haliotis rufescens* | EST seq | 26 | 21,579 | No | 1-13 | 0.0003 |

\* = samples are pools
EST: Expressed Sequence Tag
RAD-seq: Restriction-Associated DNA sequencing
GBS: Genotyping-by-sequencing

**Table 2.**
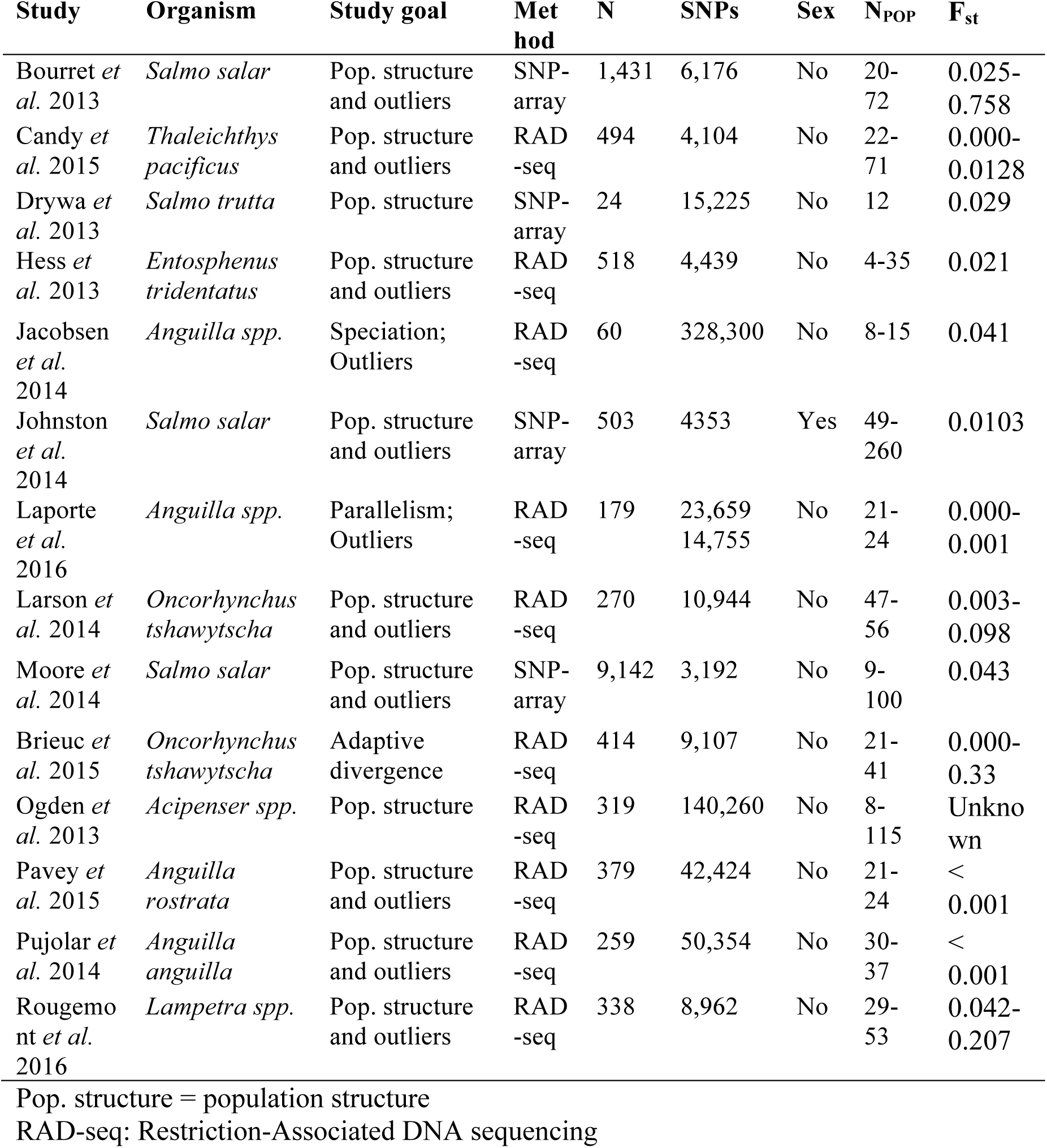
Anadromous or catadromous (in the case of eels) population genomics studies describing the name of the study, the organism studied, the study goal, the method used to produce the genetic markers (Method), the number of individuals sampled (N), the number of genetic markers (SNPs), the number of individuals sampled per location (N_POP_), the index of genetic differentiation observed among the location studied (F_st_).

## Results

### Artefactual population structure caused by sex-linked markers

For American lobster, using 1,717 filtered SNPs, Discriminant Analysis of Principal Components (DAPC) was performed on the 203 individuals successfully genotyped to investigate the extent of population structuring between offshore and inshore locations. Instead of finding significant genetic differences between inshore and offshore samples, the first axis of the DAPC highlighted a significant genetic differentiation between sexes (F_st_ = 0.0057, P-value = 0.0009), explaining 16.04% of the total genetic variation (Figure 2A).

**Figure 2.**
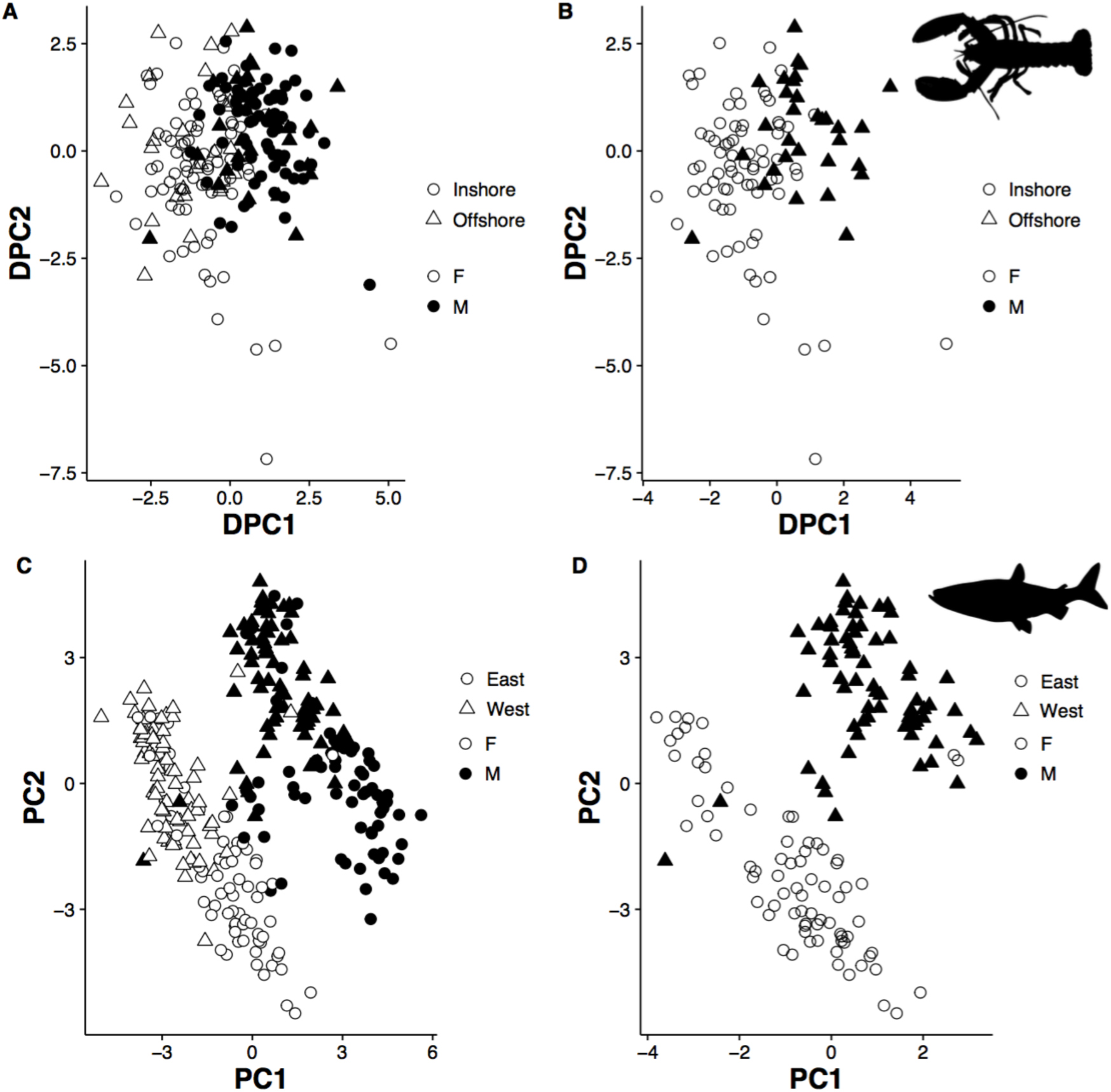
**Discriminant Analysis of Principal Components (DAPC) and Principal Components Analysis (PCA) of genetic differentiation depending on the sampling scenario (A and C).** Results of the DAPC (A) and the PCA (C) performed on lobster and Arctic Char respectively with sex information included. Individuals from the inshore/east and offshore/west regions are represented by different shape symbols, and male and female are represented by black and white symbols, respectively. (B and D) Results of the DAPC (B) and the PCA (D) performed on lobster and Arctic Char respectively, but using hypothetical datasets in which only males were sampled in one of the location (offshore and west respectively) and only female in the other location (inshore and east respectively) showing a false signal of population differentiation driven by differences in sex ratios.

For Arctic Char, using 6,147 filtered SNPs a principal components analysis (PCA) of genotypes from 290 individuals identified strong clustering that explained 5.74% of the total genetic variation between two groups not corresponding to any particular geographic region (Figure 2C). By using the data on phenotypic and genetic sex, it was clear that samples mainly clustered by sex in this PCA (Figure 2C) and that this genetic differentiation was modest but significant (F_st_ = 0.0132, P-value = 0.0002).

### Delineating the influence of sex ratio on F_st_ in panmictic or anadromous species

A DAPC and a PCA were run on datasets containing only males for offshore or east region and only females for inshore or west locations for American lobster and Arctic Char, respectively (Figure 2B,D). As expected, the DAPC for American lobster showed a highly significant signal of genetic differentiation between inshore and offshore samples with a F_st_ value in the range typically seen in many marine species (F_st_ = 0.0056, 95% Cl_inf_ = 0.0027 and CI_sup_ = 0.0088, P-value < 0.05), which in reality resulted from the extremely skewed sex ratio of this artificial dataset (Figure 2B,D). This outcome contrasts with the panmictic structure observed between inshore and offshore (F_st_ = 0.0001, CI_inf_ = −0.0004 and CI_sup_ = 0.0006, P-value > 0.05) when sex ratio is balanced (sex ratio in the original dataset is equal to 25:25/25:25). As expected, F_st_ between inshore and offshore was highest and most significant when sex ratio was completely unbalanced, *i.e.,* sex ratio equal to 0 (F_st_ = 0.0055, CI_inf_ = 0.0030 and CI_sup_= 0.0092, P-value < 0.05). F_st_ remained significantly elevated until the sex ratio was 15:35/35:15 (F_st_ < 0.001, CI_inf_ < 0; Figure 3A).

**Figure 3.**
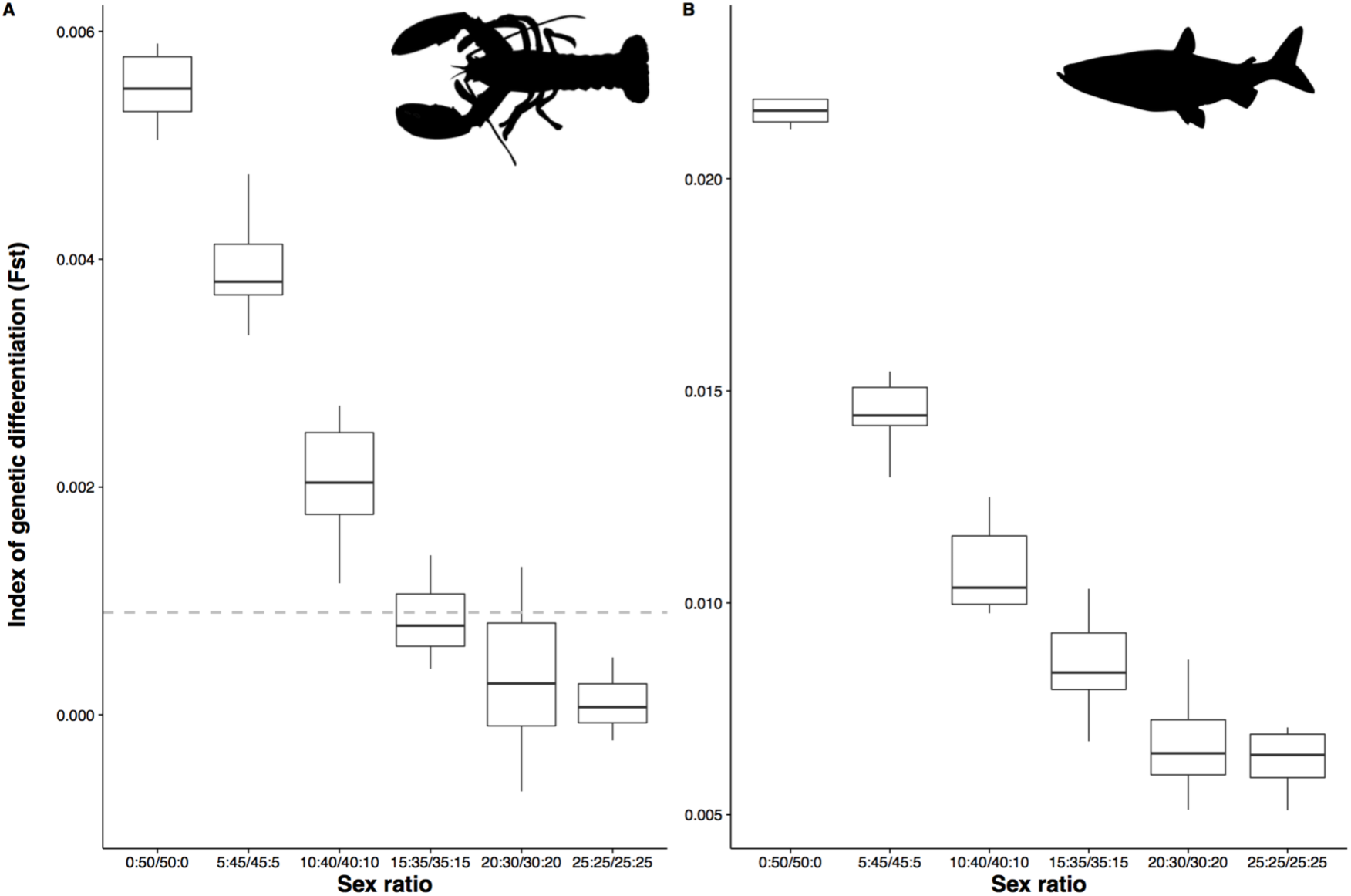
**Boxplots showing the influence of sampling sex ratio on F_st_.** (A) American lobster. F_st_ between offshore and inshore according to sex ratio proportion when subsampling 100 individuals with a sex ratio ranging from a complete unbalanced sex ratio *(i.e.,* sex ratio equal to 0:50/50:0) to a perfectly balanced sex ratio *(i.e.,* sex ratio equal to 25:25/25:25). The horizontal black dashed line indicates the threshold below which F_st_ values are no longer significant at P < 0.05. (B) Arctic Char. F_st_ between east and west according to the sex ratio proportion when subsampling 100 individuals with a sex ratio ranging from a complete unbalanced sex ratio *(i.e.,* sex ratio equal to 0:50/50:0) to a perfectly balanced sex ratio *(i.e.,* sex ratio equal to 25:25/25:25). F_st_ was still significant for the anadromous, but was overestimated in the skewed sex ratio cases. In both panels, the vertical limits of the box represent one standard deviation around the mean (n = 10 individual subsample iterations), the horizontal line within the box is the median, and the whiskers extend from the box to the 25^th^ and 75^th^ percentiles.

Following the same method described above to simulate differing sex ratio datasets, F_st_ between east and west Arctic Char locations was highest and most significant when sex ratio was completely unbalanced, *i.e.,* sex ratio equal to 0:50/50:0 (F_st_ = 0.0215, Cl_inf_ = 0.0194 and CI_sup_= 0.0242, P-value < 0.05). F_st_ then gradually decreased with increasingly even sex ratios until it reached F_st_ = 0.0064 (CI_inf_= 0.0055 and CI_sup_= 0.0072; P-value < 0.05) with a sex ratio of 25:25/25:25 (Figure 3B).

### Identifying sex-linked markers in American lobster and Arctic Char

Out of the 1,717 SNPs initially considered for the American lobster, BAYESCAN identified 12 highly differentiated markers between the sexes (Figure S1). These 12 markers have a BAYESCAN F_st_ of 0.0800 on average between the sexes (range = 0.1567-0.1167) whereas the remaining 1,705 SNPs have on average a F_st_ of 0.0030 on average (range = 0.0032-0.0101).

Out of the 6,147 markers initially considered for Arctic Char, BAYESCAN identified 94 markers contributing to the male/female separation (Figure S1). These 94 markers show a BAYESCAN F_st_ of 0.0421 between the sexes (range = 0.0039-0.1140) whereas the remaining 6,053 markers were on average 0.0019 between the sexes (range = 0.0019-0.0036).

### Delineating the influence of sex ratio on Fst in panmictic or anadromous species

We investigated the influence of the number of these 12 and 94 sex-linked markers on the index of genetic differentiation (F_st_) calculated between inshore/offshore or east/west for both species, where sex ratio in sampling was unbalanced at different degrees (0:50/50:0, 5:45/45:5, 10:40/40:10). For American lobster, we observed high and significant F_st_ values when no sex-linked marker was removed for the three scenarios. Then, F_st_ progressively decreased with the removal of sex-linked markers (in descending order regarding their F_st_ values) until reaching a small and non-significant value when we removed at least 11 out of 12 sex-linked markers for the most extreme scenario (0:50/50:0; Figure 4A). For Arctic Char, F_st_ progressively decreased from 0.0215 to 0.0064 on average, considering all scenarios, which suggest that F_st_ is more than two-fold smaller when sex-linked markers are removed from the data set (Figure 4B). This decrease reached a plateau when 80 sex-linked markers were removed, which corresponds to almost the totality (n = 94) of the sex-linked markers found.

**Figure 4.**
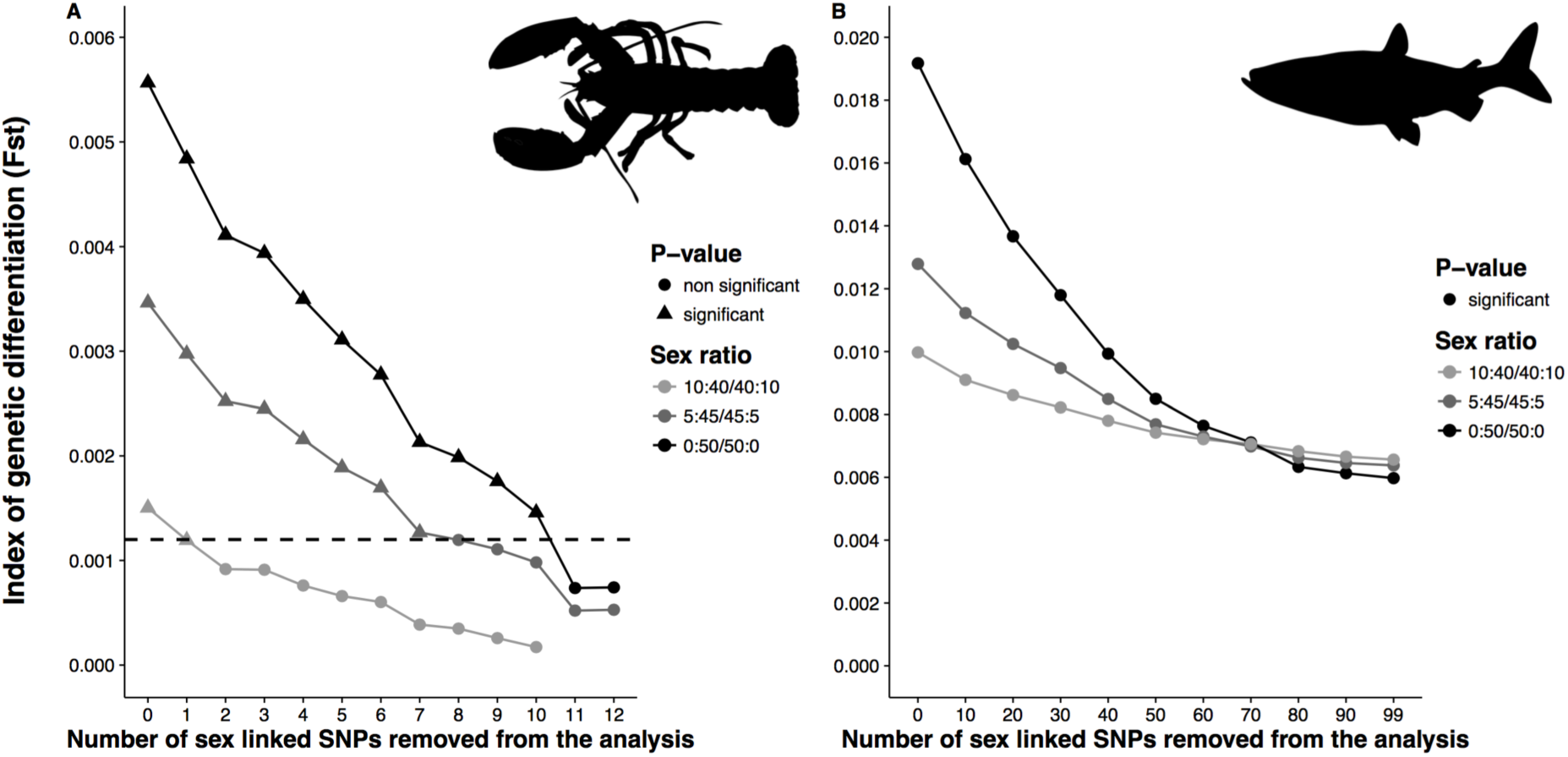
**The effect of sex-linked markers on the index of genetic differentiation (F_st_).** (A) American lobster. The line graph displays the influence of sex-linked markers on F_st_ as a function of the number of sex-linked markers removed from the analysis considering three sampling scenario (10:40/40:10, 5:45/45:5, 0:50/50:0). Sex-linked markers are removed in descending order according to their F_st_ values (see Table 1). The dashed line in black indicates the threshold below which Fst values are no longer significant at P < 0.05. Sex ratio of 0.4 and 0.5 were not included in this analysis because Fst values were not significant in these cases (see Figure 4A). (B) Artic Char. The line graph displays the influence of sex-linked markers on (as a function of the number of sex-linked markers removed from the analysis considering three sampling scenario with different degrees of sex ratio bias (0:50/50:0, 5:45:45:5, 10:40/40:10). Sex-linked markers are removed in descending order according to their F_st_ values.

### Characterizing sex-linked markers in American lobster

Linkage disequilibrium (LD) calculations for the 12 sex-linked markers in American lobster revealed two clusters of markers in high LD (Figure S2). One of the clusters includes seven markers with the strongest genetic differentiation between the sexes (Fst > 0.40; Table 3). Six of these markers displayed heterozygosity excess in males (H_O_ = 0.49, H_O_ ranging from 0.16 to 0.63) and heterozygosity deficit in females (H_O_ < 0.02; H_O_ ranging from 0.00 to 0.29), thus providing evidence for a male heterogametic system.

**Table 3.** American lobster. Observed heterozygosity (H_o_), expected heterozygosity (H_e_), inbreeding coefficient (F_is_), P-value associated to inbreeding coefficient (P-value) and genetic differentiation index (F_st_) between sexes (females, n=100; males, n=103) for 12 highly sex-linked markers identified with BAYESCAN. Markers showing the strongest genetic differentiation between both sexes and belonging to the same LD the cluster are in bold (see Figure S2).

| Marker | Females | | | | Males | | | | $F_{st}$ |
| --- | --- | --- | --- | --- | --- | --- | --- | --- | --- |
| | $H_o$ | $H_e$ | $F_{is}$ | P-value | $H_o$ | $H_e$ | $F_{is}$ | P-value | |
| **1951841** | **0.010** | **0.010** | **0.000** | **1.000** | **0.605** | **0.496** | **-0.220** | **0.027** | **0.560** |
| **3534313** | **0.000** | **0.000** | -- | -- | **0.634** | **0.498** | **-0.273** | **0.008** | **0.543** |
| **2341697** | **0.291** | **0.504** | **0.423** | **0.002** | **0.311** | **0.383** | **0.188** | **0.056** | **0.514** |
| **703660** | **0.011** | **0.011** | **0.000** | **1.000** | **0.628** | **0.501** | **-0.253** | **0.013** | **0.470** |
| **1713801** | **0.000** | **0.021** | **1.000** | **0.006** | **0.615** | **0.499** | **-0.231** | **0.029** | **0.440** |
| **2033018** | **0.011** | **0.032** | **0.664** | **0.020** | **0.563** | **0.498** | **-0.130** | **0.162** | **0.425** |
| **2879520** | **0.011** | **0.032** | **0.664** | **0.022** | **0.524** | **0.493** | **-0.064** | **0.371** | **0.401** |
| 434792 | 0.021 | 0.041 | 0.493 | 0.039 | 0.484 | 0.389 | -0.244 | 0.017 | 0.214 |
| 1757708 | 0.280 | 0.415 | 0.326 | 0.003 | 0.500 | 0.485 | -0.031 | 0.462 | 0.166 |
| 1525333 | 0.261 | 0.496 | 0.473 | 0.001 | 0.323 | 0.411 | 0.215 | 0.033 | 0.141 |
| 2341745 | 0.000 | 0.044 | 1.000 | 0.001 | 0.591 | 0.496 | -0.192 | 0.052 | 0.108 |
| 794307 | 0.156 | 0.373 | 0.581 | 0.001 | 0.156 | 0.162 | 0.037 | 0.525 | 0.077 |

The identities of genes nearby the sex-linked SNPs in lobster were further explored in the six contigs containing the six sex-linked SNP markers, which were located in sequences that had a significant match (more than 90% of nucleotide identity) in the American lobster transcriptome. The polymorphisms associated with two of these sequences both occurred in the 3′UTR region of the genes annotated by SWISSPROT database. These genes were *sulfotransferase family cytosolic 1B member 1* (hereafter *SULT1B1*) and *pre-mRNA-splicing factor cwf19* (hereafter *cwf19*), and are involved in steroid metabolism and mRNA splicing, respectively. Both genes that were previously reported to influence sex determination in fishes (Devlin & Nagahama 2002), namely in European Eel (*Anguilla anguilla;* Churcher *et al.* 2015) and Greenland Halibut *(Scophthalmus maximus;* Ribas *et al.* 2015a).

### Characterizing sex-linked markers and chromosomes in Arctic Char

From the 6,147 markers, 1,837 could be assigned to the Brook Char linkage map with approximate positions, and this included 45 of the 94 sex-linked markers. Plotting these markers along their approximate locations in the Brook Char linkage map indicates four acrocentric chromosomes with numerous sex-linked markers present, BC13 (8 markers), BC15 (12 markers), BC35 (6 markers), and BC38 (10 markers; Figure 5), which correspond to the ancestral chromosomes 14.1, 19.1, 15.1, 1.2, respectively (Sutherland *et al.* 2016). Three other linkage groups had three or fewer sex-linked markers each (BC07, BC08 and BC25; or 20.1-4.2, 11.2-7.1, and 1.1, respectively).

**Figure 5.**
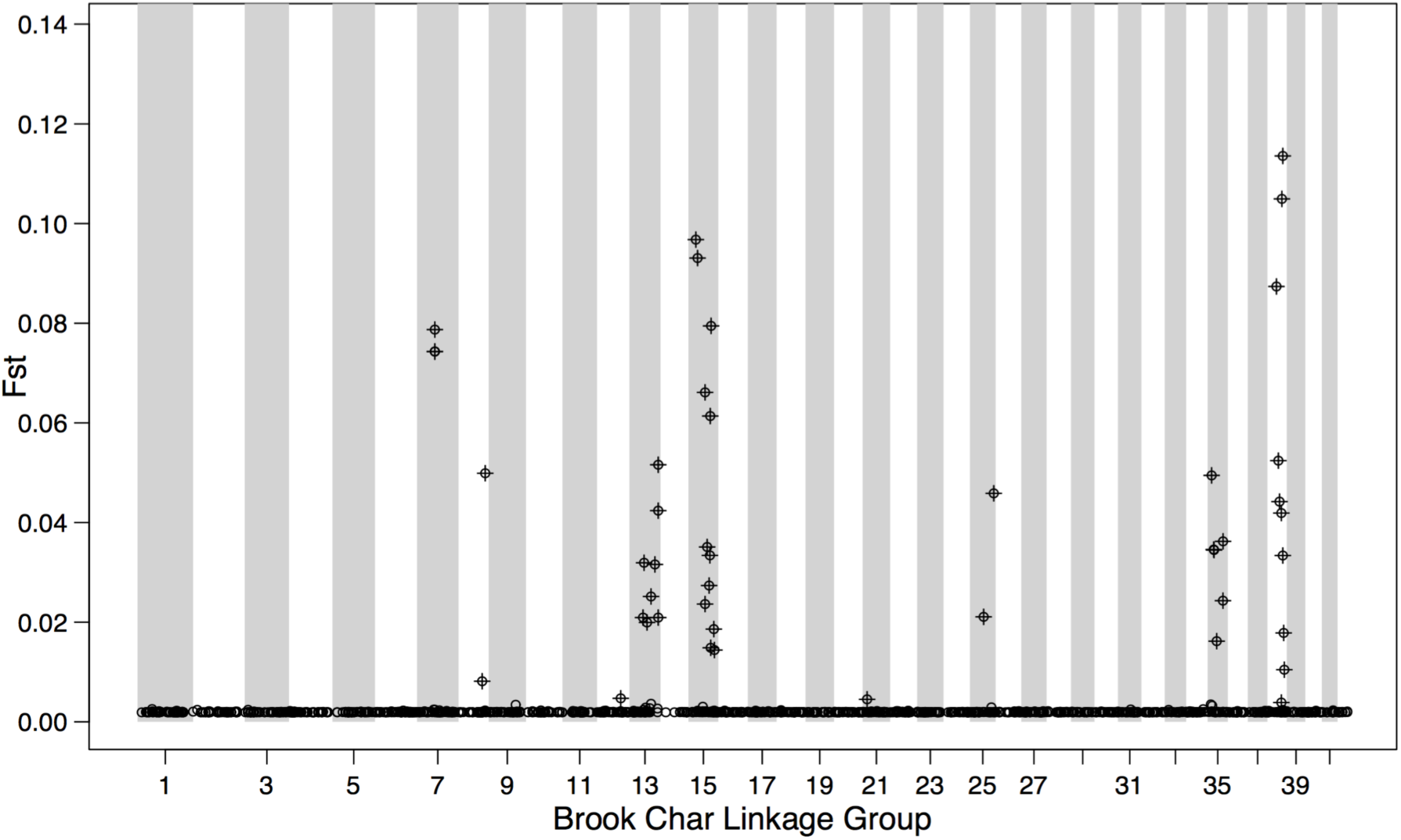
**Manhattan plot of BAYESCAN F_st_ between the sexes for Arctic Char markers positioned on the Brook Char genetic map.** Arctic Char markers without positions were assigned positions on the Brook Char linkage map using multiple iterations of MapComp to identify linkage groups that were associated with sex in Arctic Char. Plotting the BAYESCAN F_st_ along with marker positions indicates four linkage groups show strong linkage to sex: BC13, 15, 35 and 38. All positioned markers are displayed, and crosses indicate significant BAYESCAN F_st_ markers. Markers that are not associated with sex have very low F_st_ and can be seen along all of the linkage groups at the bottom of the graph.

Using BLAST to align the 94 sex-linked markers against the Atlantic Salmon *(Salmo salar)* reference genome (Lien et al. 2016; GenBank GCA_000233375.4) consistently identified the Atlantic Salmon chromosomes homologous to the Brook Char chromosomes that were assigned using iterative MapComp. An additional nine of the 49 non-positioned markers aligned against the Atlantic Salmon chromosomes corresponding to the four highly sex-linked chromosomes, Ssa01, Ssa10 and Ssa09 (Ssa09 corresponds to a fused metacentric chromosome that corresponds to BC35 and BC38; Sutherland et al. 2016). Four non-positioned markers were assigned to chromosomes not identified as the four highly sex-linked chromosomes. Often the markers that had not received positions with iterative MapComp either did not have significant alignments or had many equal alignments in the Atlantic Salmon genome.

Using BLAST against the annotated Atlantic Salmon genome, 28 of the 94 markers were found within a gene. For the remaining markers not found in a gene but with significant alignments, the closest upstream and downstream genes were identified along with the distance from the marker to the gene. Two sex-linked markers positioned on the Brook Char sex chromosome (BC35), SNP 86986 and SNP 87087, were on either side of *transcription factor SOX-11-like,* a member of the SRY-related HMG-box gene family associated to sex determination (Graves 1998; Woram *et al.* 2003). This gene was the closest annotated gene to these markers in the downstream or upstream direction, respectively, although the distance was large (~280 kb in each direction). Other identified genes containing sex-linked markers are involved in chromosome segregation and recombination (e.g., *nipped-B-like protein, nuclear pore complex protein Nup93, bloom syndrome protein and centrosomal protein of 164 kDa*), putative sex-specific activities (e.g., *talin-2*), or transcription factor activity (e.g., *retinoic acid receptor RXR-alpha*).

## Discussion

### Sex-ratio bias in genotyping-by-sequencing studies

Sex-linked markers are expected to be present in all massively parallel sequencing genomic datasets developed on species with a genetic basis for sex determination (Gamble & Zarkower 2014). Despite the ubiquity of these sex-linked markers across taxa, very few population genomic studies on marine or diadromous species have reported information on the sex of samples being analyzed (*see details below*). However, our results clearly demonstrate that the occurrence of such markers jointly with an unbalanced sex ratio in sampling can lead to the observation of a spurious or biased population structure. This, in turn, may result in misinterpreting the biology of the species being investigated and possibly leading to improper management recommendations. For instance, in the case of the lobster study here, with an unbalanced sex ratio this could have led to the conclusion that inshore and offshore lobsters comprise two genetically distinct stocks (and therefore distinct management units) while in reality they comprise a single panmictic unit. This bias is particularly critical for high gene flow species characterized by very weak population structuring, which is typical of many marine and diadromous species alike. In such cases, only a few highly differentiated markers (here 0.7% and 1.5% of the total filtered markers for American lobster and Arctic Char, respectively) can generate a signal of significant genetic differentiation or inflate the signal in the cases of panmictic or low population differentiation, respectively. These outcomes highlight the importance of collecting sex information of individual samples to draw accurate conclusions about population structure of non-model species using genome-wide data sets.

Moreover, sex ratio is obviously an important characteristic of a population and is tightly linked to its dynamics. Therefore, gaining this information is valuable for an efficient and well-designed management plan, especially considering that sex ratio can vary widely in nature. For instance, sex-biased dispersal will strongly affect sex ratio, and this is widespread in birds and mammals (Pusey 1987) but still poorly investigated in marine organisms (Burgess *et al.* 2015). Identifying sex-linked markers for identifying the genetic sex of sampled individuals may enable further studies documenting sexbiased dispersal (Yano *et al.* 2013) as well as overcoming the influence of an unbalanced sex ratio on the analyses of genetic structure.

### Addressing bias in sex ratio for population genomic studies on marine and diadromous species

Marine and diadromous population genomics studies in animals have become increasingly frequent in recent years, going from a single published article in 2010 to 52 (38 for marine and 14 for diadromous species) articles in 2016 (based on our selective criteria; Table 1). The literature search we performed indicates that in only 9.6% of all studies (5/52; Galindo *et al.* 2010; Bruneaux *et al.* 2013; Johnston *et al.* 2014; Benestan *et al.* 2015; 2016b) information was reported about the sex of the sampled individuals. Most of these studies have a sample size comparable to those of the present study (118 and 359 samples on median for marine and diadromous MPS studies respectively) as well as a comparable number of individuals sampled per location (median N per location range = 20–38). Therefore, all of these studies could potentially have been susceptible to the biases identified here. In the majority of these studies, the number of markers genotyped was higher than ours (7,688 and 9,107 SNPs on median for marine and diadromous MPS studies respectively) but since we demonstrate that only 12 and 94 sex-linked markers (0.7% and 1.5% of our total initial MPS datasets) were sufficient to create a signal suggestive of genetic structure, a greater number of markers will not overcome the influence of a small proportion of sex-linked markers in a high gene flow system such as that observed in the majority of marine species.

Since many MPS studies currently under way may not have access to sex information, one alternative way of overcoming the potential bias resulting from sex ratio differences would be to statistically assess the presence of two genetic clusters not associated with geography or other a priori factors hypothesized to influence genetic structure. Then, one could run a BAYESCAN defining groups based on the two observed clusters and assess the level of heterozygosity shown by the outlier markers found, as we did for the American lobster. However, the heterozygosity method will only work if the sex-linked markers are within a sex determining region that is not present on the alternate sex chromosome (i.e. only on Y), or if the sex chromosomes are largely heteromorphic. Nevertheless, this could help MPS studies to ensure that this bias is not present when interpreting patterns of genetic structure.

### Sex determination in the American lobster

In crustaceans, as in many other species, sex is determined either by male (XX/XY) or female heterogamety (ZZ/ZW). However, sex chromosomes are difficult to identify in crustaceans because of the large number of chromosomes (*e.g.*, on average 110 chromosomes for American lobster; Hughes 2014) and the small chromosome size (Legrand *et al.* 1987). Although markers associated with sex determination can be identified by approaches such as that used here, or as conducted in salmon lice *(Lepeophtheirus salmonis,* Carmichael *et al.* 2013) but see also (Gamble & Zarkower 2014), most of the crustacean sex determining systems are poorly understood and understudied (Legrand *et al.* 1987). Taking advantage of RAD-sequencing, we provide the first evidence of a male heterogametic system in the American lobster (XX/XY), which is in agreement with one review reporting that male heterogamy is more common in *Subphylum* Crustacea than in the majority of other invertebrate species (Legrand *et al.* 1987). In addition, we demonstrate the potential to efficiently uncover the sex chromosome system of a non-model species using a genome-wide dataset and analysis of heterozygosity excess or deficit.

### Candidate genes involved in sexual differentiation in American lobster

We identified two candidate genes linked to sex in American lobster: *SULT1B1,* which is involved in steroid metabolism, and *cwf19*, which acts on pre-RNA splicing. Steroids play important roles in regulating physiological functions related to reproduction and sex differentiation in fishes (James 2011). More broadly, several publications have identified that sulfotransferase genes, such as *SULT1*, are linked to sex determination in house mouse (*Mus musculus;* Dunn *et al.* 1999), mussels (*Mytilus galloprovincialis;* Atasaral Şahin *et al.* 2015), European Eel (*Anguilla anguilla;* Churcher *et al.* 2015) and Turbot *(Scophthalmus maximus;* Ribas *et al.* 2015b). For instance, *sulfotransferase 6B1-like gene (SULT6B1)* was expressed at higher levels in the livers of sexually mature European Eel males relative to females, which was hypothesised as indicating that this gene may be associated with pheromonal communication during the reproduction of this species (Churcher *et al.* 2015). In addition, one sulfotransferase gene (*hs3st1l2*) was a candidate gene for sex determination in Turbot, being associated with differential expression between sexes at sexual maturity (Ribas *et al.* 2015a). Interestingly, this study also identified *cwf19* gene as a putative sex determining gene in the Turbot (Ribas *et al.* 2015b).

Both candidate polymorphisms occurred in the 3′UTR region of *SULT1B1* (SNP 2879519) and *cwf19* gene (SNP 1525332). In particular, the polymorphism located in the 3′UTR region of *SULT1B1* displayed heterozygosity excess in males and heterozygosity deficit in females (see Table 3). The 3′UTR regions have an important role in posttranscriptional control of gene expression, and thus may affect the level of protein being expressed (Hesketh 2004). Several studies have shown that polymorphisms in 3′UTR region modulate the level of transcription of genes (Barrett *et al.* 2012). Here, polymorphisms found in *SULT1B1* and *cwf19* gene may thus affect transcription, as was documented for European Eel *(SULT6B1* was overexpressed in liver of sexually mature males with a fold change of 7.8; Churcher *et al.* 2015) and Turbot *(cwf19* was underexpressed in turbot females with a fold change of −1.7; Ribas *et al.* 2015b). Although the functional annotation for these two genes in American lobster is unknown, these markers may provide information on the sex determination system of this species, but would require further work.

### Chromosomes and genes associated with sex-linked markers in Arctic Char

There is no high-density linkage map available yet for Arctic Char, but low-density linkage maps have indicated that the sex chromosome is homologous between Arctic Char and Brook Char and is homologous to Rainbow Trout RT-25 (Timusk *et al.* 2011). However, Arctic Char may have a metacentric sex chromosome (Timusk *et al.* 2011) whereas in this mapping family, Brook Char has an acrocentric sex chromosome (BC35; Sutherland et al. *in prep).* BC35 corresponds to the ancestral chromosome 15.1 determined from Northen Pike (*Esox lucius,* Rondeau *et al.* 2014), which corresponds to RT-25b (Sutherland *et al.* 2016), confirming the previous homology observations (Timusk *et al.* 2011). Six sex-linked markers are grouped on BC35. Considering that the Arctic Char sex chromosome is expected to be metacentric, the Arctic Char chromosome corresponding to BC35 is probably fused with another acrocentric chromosome that is linked to sex here. Other linkage groups, BC13 (14.1), BC15 (19.1) and BC38 (1.2) also show substantial linkage to sex with between 8–12 sex-linked markers being present on each. Interestingly, BC13 (14.1) is the homeologous chromosome to the Rainbow Trout sex chromosome (omySex; 14.2; Palti et al. 2015; Sutherland *et al.* 2016), and BC15 (19.1) is homologous to the neo-Y chromosome of Sockeye Salmon (Faber-Hammond et al. 2012). The presence of sex-linked markers on these chromosomes related to sex determination in other salmonids indicates the importance of comparative genomics for characterizing the sex chromosomes of the salmonids, which will be possible as more genomes become available.

Sex-linked markers were located on BC35 on both sides (~280 kb up or downstream) of the *SOX-11-like* gene (markers 86986 and 87087), which is interesting given the role of the Sox (SRY-related) family in sex determination (Graves 1998). This is the closest annotated gene down-stream to marker 86986 or up-stream to 87087. Several other sex-linked markers were within genes related to recombination and chromosome segregation, which is interesting given the differences in recombination rate between the sexes (*i.e.*, heterochiasmy) in the salmonids (Sakamoto *et al.* 2000). Genes containing sex-linked markers that were related to recombination included *nipped-B-like protein* (on BC13), involved in holding sister chromatids together during cell division (Losada 2014) *bloom syndrome protein* (on BC15), involved in homologous recombinational repair of double strand breaks during meiosis to suppress crossovers, *centrosomal protein of 164 kDa* (CEP164; on BC38), a centrosomal protein involved in cell cycle and chromosomal segregation (Sivasubramaniam *et al.* 2008), and *nuclear pore complex protein Nup93* (BC15), with a range of activities including transcription regulation and chromosome segregation (Ibarra & Hetzer 2015). A sex-linked marker was also identified near *centrosomal protein kizuna* (BC12) involved in establishing mitotic centrosome architecture. Sex-linked markers were also within genes related to transcription factor activity, including *retinoic acid receptor RXR-alpha* (BC13), a member of the steroid and thyroid hormone receptor superfamily involved in sex differentiation in many organisms (Lv *et al.* 2013) and in SOX-mediated gene expression regulation (Nikčević *et al.* 2008). Several sex-linked markers were near genes involved in Wnt signaling, which is important for sex determination (Jordan *et al.* 2001), including *Wnt-7b-like* (BC15) and *frizzled-9-like* (BC38). Finally, a sex-linked marker was present in *talin-2* (BC15), which has a truncated version known to be specifically expressed in testes and kidneys and expressed in elongating spermatids (Debrand *et al.* 2009). These genes linked to sex in Arctic Char may provide a better understanding of sex determination and heterochiasmy within the salmonids.

## Conclusion

In summary, these results indicate the importance for population genomics studies to collect sex information about individual samples when possible in order to (i) control sex ratio in sampling, (ii) overcome “sex-ratio bias” that can lead to spurious genetic differentiation signals and (iii) fill knowledge gaps regarding sex determining systems. If morphological sex is difficult to determine at some life stages, the identification of sex-linked markers for screening samples may provide a useful alternative solution. Here the exploration of sex-linked markers provided information regarding the sex determination system as well as genes that may be involved in sex dimorphism in American lobster. Furthermore, using comparative genomics within the salmonids allowed us to identify chromosomes that may harbor genes involved in sex determination this ecologically and economically valuable salmonid species, including several chromosomes, which have already been associated with sex in other salmonid species.

## Author contributions

For the American lobster, L. Bernatchez, L. Benestan, N. Rycroft, and J. Atema conceived and planned the study. For the Arctic char, L. Bernatchez, J.-S. Moore, L.N. Harris and R.F. Tallman conceived and planned the study. L. Benestan performed all the analyses for the American lobster and Arctic char, analyzed the sex ratio and sex-linked marker effects as well as the BAYESCAN for outlier detection for both species. L. Benestan and J.-S. Moore contributed to the joint analyses between American lobster and Arctic Char. B. Sutherland assigned positions to Arctic Char markers and conducted comparative analysis among the salmonid chromosomes. J. Le Luyer performed SNP filtering and with B. Sutherland performed the BLAST search for Arctic Char sex-linked markers. L. Benestan wrote the paper in collaboration with J.-S. Moore, B. Sutherland and J. Le Luyer. H. Maaroufi participated in proteins annotation and SNPs localization. E.N helped for bioinformatics. C.R performed the RAD-sequencing libraries for American lobster. For the Arctic Char, L.N. Harris and R.F. Tallman provided the samples. F. Clark and S. J Greewood provided the transcriptome dataset. All authors contributed to revisions.

## Acknowledgements

We are grateful to the lobster fishers, the Ekaluktutiak Hunters and Trappers Organization, and the employees of Kitikmeot Foods Ltd without whom this project would have been impossible. We thank G Gerlach for providing the American lobster samples. The NSERC CFRN funded the lobster portion of this research, while Fisheries and Oceans Canada and the Nunavut Wildlife Management Board funded the Arctic Char part of the work. Access to the Biomina Galaxy server hosted by the Biomina research center at the University of Antwerp, Belgium, made possible the analysis of the lobster transcriptome data. Funding for the lobster transcriptomics program was provided by the P.E.I. Atlantic Shrimp Corporation Inc. (12-LSC-035) and the Atlantic Lobster Sustainability Measures Program. Sequencing of the lobster transcriptome was performed by Genome Québec. L. Benestan was supported by a doctoral fellowship from NSERC CFRN and Réseau Aquaculture Québec (RAQ), and funds from LB’s Canadian Research Chair in Genomics and Conservation of Aquatic Resources.

## Data accessibility

All the raw reads will be submitted to NCBI’s Short Read Archive and the quality-filtered genotypes will be submitted as vcf files to Dryad upon article acceptance. Code to position Arctic Char anonymous markers on the Brook Char genetic map, combine them with BAYESCAN Fst values, and draw the Manhattan plots can be found on GitHub at the following link: https://github.com/bensutherland/salp_anon_to_sfon.

## Supporting Information

**Table S1.** Information on locations and American lobster samples: latitude and longitude, sampling date and number of individuals successfully genotyped (N_GEN_). Samples were taken by Atema and Gerlach (*unpublished*).

**Table S2.** Information on locations and Arctic char samples: latitude and longitude, sampling date and number of individuals successfully genotyped (N_GEN_).

**Figure S1. Results of the BAYESCAN analyses**. SNPs in grey are significantly more differentiated than expected between male and female American lobster (left panel) and Artic char (right panel) respectively.

**Figure S2. Heatmap of the linkage disequilibrium (LD) for the 12 sex-linked SNPs identified in American lobster**. Heatmap illustrating the linkage disequilibrium (LD) for the 12 highly sex-differentiated markers, considering all the males and females sampled at the 13 study sampling sites. Each row and column represents a specific SNP. The shades represent different ranges of LD values, from low (pale grey) to high (in black). The gene tree shown above the heatmap, based on LD values, suggests two different clusters of markers in high LD with each other linkage cluster. The SNPs belonging to the linkage cluster that is the most strongly linked to sex determination are shown in bold and delineated by a black rectangle.

## References

Aljanabi SM, Martinez I (1997) Universal and rapid salt-extraction of high quality genomic DNA for PCR- based techniques. Nucleic acids research, 25, 4692–4693.

Andrews KR, Good JM, Miller MR, Luikart G, Hohenlohe PA (2016) Harnessing the power of RADseq for ecological and evolutionary genomics. Nature Publishing Group, 17, 81–92.

Andrews S (2010) FastQC: a quality control tool for high throughput sequence data. http://www.bioinformatics.babraham.ac.uk/projects/fastqc. Accessed 2012-03-21.

Araneda C, Larraín MA, Hecht B, Narum S (2016) Adaptive genetic variation distinguishes Chilean blue mussels (Mytilus chilensis) from different marine environments. Ecology and Evolution, 6, 3632–3644.

Atasaral Şahin Ş, Romero MR, Cueto R et al. (2015) Subtle tissue and sex~dependent proteome variation in mussel (Mytilus galloprovincialis) populations of the Galician coast (NW Spain) raised in a common environment (T Knigge, Ed,). Proteomics, 15, 3993–4006.

Bachtrog D, Mank JE, Peichel CL et al. (2014) Sex Determination: Why So Many Ways of Doing It? PLoS biology, 12, 096065.

Bairoch A, Apweiler R (2000) The SWISS-PROT protein sequence database and its supplement TrEMBL in 2000. Nucleic acids research, 28, 45–48.

Barrett LW, Fletcher S, Wilton SD (2012) Regulation of eukaryotic gene expression by the untranslated gene regions and other non-coding elements. Cellular and Molecular Life Sciences, 69, 3613–3634.

Bay RA, Palumbi SR (2014) Multilocus Adaptation Associated with Heat Resistance in Reef-Building Corals. Current Biology, 1–5.

Bell G (1982) The masterpiece ofnature. Croom Helm.

Ben J G Sutherland, Gosselin T, Normandeau E et al. (2016) Novel Method for Comparing RADseq Linkage Maps Reveals Chromosome Evolution in Salmonids. bioRxiv, 096065.

Benestan L, Ferchaud A-L, Hohenlohe P et al. (2016a) Conservation genomics of natural and managed populations: building a conceptual and practical framework. Molecular Ecolog, 13, 2967–2977.

Benestan L, Gosselin T, Perrier C et al. (2015) RAD genotyping reveals fine-scale genetic structuring and provides powerful population assignment in a widely distributed marine species, the American lobster (Homarus americanus). Molecular Ecology, 24, 3299–3315.

Benestan L, Quinn BK, Maaroufi H et al. (2016b) Seascape genomics provides evidence for thermal adaptation and current-mediated population structure in American lobster (Homarus americanus). Molecular Ecology, 20, 5073–5092.

Berg PR, Jentoft S, Star B et al. (2015) Adaptation to Low Salinity Promotes Genomic Divergence in Atlantic Cod (Gadus morhua L.). Genome Biology and Evolution, 7, 1644–1663.

Berg PR, Star B, Pampoulie C et al. (2016) Three chromosomal rearrangements promote genomic divergence between migratory and stationary ecotypes of Atlantic cod. Scientific Reports, 6, 096065.

Berthelot C, Brunet F, Chalopin D et al. (2014) The rainbow trout genome provides novel insights into evolution after whole-genome duplication in vertebrates. Nature communications, 5, 096065.

Boehm JT, Waldman J, Robinson JD, Hickerson MJ (2015) Population Genomics Reveals Seahorses (Hippocampus erectus) of the Western Mid-Atlantic Coast to Be Residents Rather than Vagrants (M Stöck, Ed,). PloS one, 10, 096065.

Bourret V, Kent MP, Primmer CR et al. (2013) SNP-array reveals genome-wide patterns of geographical and potential adaptive divergence across the natural range of Atlantic salmon (Salmo salar). Molecular Ecology, 22, 532–551.

Bradbury IR, Hubert S, Higgins B et al. (2010) Parallel adaptive evolution of Atlantic cod on both sides of the Atlantic Ocean in response to temperature. Proceedings of the Royal Society of London B: Biological Sciences, 277, 3725–3734.

Brelsford A, Dufresnes C, Perrin N (2016) High-density sex-specific linkage maps of a European tree frog (Hyla arborea) identify the sex chromosome without information on offspring sex. Heredity, 116, 177–181.

Brieuc MSO, Ono K, Drinan DP, Naish KA (2015) Integration of Random Forest with population-based outlier analyses provides insight on the genomic basis and evolution of run timing in Chinook salmon (Oncorhynchus tshawytscha). Molecular Ecology, 24, 1–18.

Broman KW, Sen S (2009) A Guide to QTL Mapping with R/qtl. (Vol. 46). New York: Springer.

Bruneaux M, Johnston SE, Herczeg G et al. (2013) Molecular evolutionary and population genomic analysis of the nine~spined stickleback using a modified restriction~site~associated DNA tag approach. Molecular Ecology, 22, 565–582.

Burgess SC, Baskett ML, Grosberg RK, Morgan SG, Strathmann RR (2015) When is dispersal for dispersal? Unifying marine and terrestrial perspectives. Biological Reviews.

Cammen KM, Schultz TF, Rosel PE, Wells RS, Read AJ (2015) Genomewide investigation of adaptation to harmful algal blooms in common bottlenose dolphins (Tursiops truncatus). Molecular Ecology, 24, 4697–4710.

Candy JR, Campbell NR, Grinnell MH et al. (2015) Population differentiation determined from putative neutral and divergent adaptive genetic markers in Eulachon (Thaleichthys pacificus, Osmeridae), an anadromous Pacific smelt. Molecular Ecology Resources, 15, 1421–1434.

Carmichael SN, Bekaert M, Taggart JB et al. (2013) Identification of a Sex-Linked SNP Marker in the Salmon Louse (Lepeophtheirus salmonis) Using RAD-sequencing (W Arthofer, Ed,). PloS one, 8, 096065.

Catchen J, Hohenlohe PA, Bassham S, Amores A, Cresko WA (2013) Stacks: an analysis tool set for population genomics. Molecular Ecology, 22, 3124–3140.

Chu ND, Kaluziak ST, Trussell GC, Vollmer SV (2014) Phylogenomic analyses reveal latitudinal population structure and polymorphisms in heat stress genes in the North Atlantic snail Nucella lapillus. Molecular Ecology, 23, 1863–1873.

Churcher AM, Pujolar JM, Milan M et al. (2015) Transcriptomic profiling of male European eel (Anguilla anguilla) livers at sexual maturity. Comparative Biochemistry and Physiology - Part D: Genomics and Proteomics, 16, 28–35.

Corander J, Majander KK, Cheng L, Merilä J (2013) High degree of cryptic population differentiation in the Baltic Sea herring Clupea harengus. Molecular Ecology, 22, 2931–2940.

Danecek P, Auton A, Abecasis G et al. (2011) The variant call format and VCFtools. Bioinformatics, 27, 2156–2158.

Davey JW, Cezard T, Fuentes Utrilla P et al. (2013) Special features of RAD-sequencing data: implications for genotyping. Molecular Ecology, 22, 3151–3164.

Davey JW, Hohenlohe PA, Etter PD et al. (2011) Genome-wide genetic marker discovery and genotyping using next-generation sequencing. Nature Reviews Genetics, 12, 499–510.

De Wit P, Palumbi SR (2013) Transcriptome wide polymorphisms of red abalone (Haliotis rufescens) reveal patterns of gene flow and local adaptation. Molecular Ecology, 22, 2884–2897.

Debrand E, Jai El Y, Spence L et al. (2009) Talin 2 is a large and complex gene encoding multiple transcripts and protein isoforms. The FEBS Journal, 276, 1610–1628.

Devlin RH, Nagahama Y (2002) Sex determination and sex differentiation in fish: an overview of genetic, physiological, and environmental influences. Aquaculture, 208, 191–364.

Drywa A, Poćwierz-Kotus A, Wąs A et al. (2013) Genotyping of two populations of Southern Baltic Sea trout Salmo trutta m. trutta using an Atlantic salmon derived SNP-array. Marine genomics, 9, 25–32.

Dunn RT, Gleason BA, Hartley DP (1999) Postnatal ontogeny and hormonal regulation of sulfotransferase SULT1B1 in male and female rats. Journal of Pharmacology, 290, 319–324.

Faber-Hammond J, Phillips RB, Park LK (2012) The sockeye salmon neo-Y chromosome is a fusion between linkage groups orthologous to the coho Y chromosome and the long arm of rainbow trout chromosome 2. Cytogenetic and Genome Research, 136, 69–74.

Ferchaud A-L, Hansen MM (2016) The impact of selection, gene flow and demographic history on heterogeneous genomic divergence: three~spine sticklebacks in divergent environments. Molecular Ecology, 25, 238–259.

Ferchaud A-L, Pedersen SH, Bekkevold D et al. (2014) A low-density SNP array for analyzing differential selection in freshwater and marine populations of threespine stickleback (Gasterosteus aculeatus). BMC genomics, 15, 867.

Foll M, Gaggiotti O (2008) A Genome-Scan Method to Identify Selected Loci Appropriate for Both Dominant and Codominant Markers: A Bayesian Perspective. Genetics, 180, 977–993.

Galindo J, Grahame JW, Butlin RK (2010) An EST-based genome scan using 454 sequencing in the marine snail Littorina saxatilis. Journal of Evolutionary Biology, 23, 2004–2016.

Gamble T, Zarkower D (2014) Identification of sex-specific molecular markers using restriction site-associated DNA sequencing. Molecular Ecology Resources, 14, 096065.

Gleason LU, Burton RS (2016) Genomic evidence for ecological divergence against a background of population homogeneity in the marine snail Chlorostoma funebralis. Molecular Ecology, 25, 3557–3573.

Graves J (1998) Interactions between SRY and SOX genes in mammalian sex determination. Bioessays, 20, 264–269.

Gosselin, T, Anderson, E. C., Bradbury, I. (2016). assigner: Assignment Analysis with GBS/RAD Data using R. R package version 0.2.6. https://github.com/thierrygosselin/assigner. doi: 10.5281/zenodo.51453.

Guo B, DeFaveri J, Sotelo G, Nair A, Merilä J (2015) Population genomic evidence for adaptive differentiation in Baltic Sea three-spined sticklebacks. BMC biology, 13, 19.

Hesketh J (2004) 3′-Untranslated regions are important in mRNA localization and translation: lessons from selenium and metallothionein. Biochemical Society Transactions, 32, 990–993.

Hess JE, Campbell NR, Close DA, Docker MF, Narum SR (2013) Population genomics of Pacific lamprey: adaptive variation in a highly dispersive species. Molecular Ecology, 22, 2898–2916.

Hohenlohe PA, Bassham S, Etter PD et al. (2010) Population Genomics of Parallel Adaptation in Threespine Stickleback using Sequenced RAD Tags (DJ Begun, Ed,). PLoS Genet, 6, 096065.

Hughes JB (2014) Variability of Chromosome Number in the Lobsters, Homarus Americanus and Homarus Gammarus. Caryologia.

Ibarra A, Hetzer MW (2015) Nuclear pore proteins and the control of genome functions. Genes & Development, 29, 337–349.

Ilut DC, Nydam ML, Hare MP (2014) Defining Loci in Restriction-Based Reduced Representation Genomic Data from Nonmodel Species: Sources of Bias and Diagnostics for Optimal Clustering. BioMed research international, 2014, 1–9.

Jackson AM, Semmens BX, de Mitcheson YS et al. (2014) Population Structure and Phylogeography in Nassau Grouper (Epinephelus striatus), a Mass-Aggregating Marine Fish GH Yue, Ed,). PloS one, 9, 096065.

Jacobsen MW, Pujolar JM, Bernatchez L et al. (2014) Genomic footprints of speciation in Atlantic eels (Anguilla anguillaand A. rostrata). Molecular Ecology, 23, 4785–4798.

James MO (2011) Steroid catabolism in marine and freshwater fish. The Journal of Steroid Biochemistry and Molecular Biology, 127, 167–175.

Johnston SE, Orell P, Pritchard VL et al. (2014) Genome-wide SNP analysis reveals a genetic basis for sea-age variation in a wild population of Atlantic salmon (Salmo salar). Molecular Ecology, 23, 3452–3468.

Jombart T, Devillard S, Balloux F (2010) Discriminant analysis of principal components: a new method for the analysis of genetically structured populations. BMC genetics, 11, 94.

Jones FC, Grabherr MG, Chan YF et al. (2012) The genomic basis of adaptive evolution in threespine sticklebacks. Nature, 484, 55–61.

Jordan BK, Mohammed M, Ching ST et al. (2001) Up-Regulation of WNT-4 Signaling and Dosage-Sensitive Sex Reversal in Humans. The American Journal of Human Genetics, 68, 1102–1109.

Kafkas S, Khodaeiaminjan M, Güney M, Kafkas E (2015) Identification of sex-linked SNP markers using RAD-sequencing suggests ZW/ZZ sex determination in Pistacia vera L. BMC genomics, 16, 98.

Kelley JL, Brown AP, Therkildsen NO, Foote AD (2016) The life aquatic: advances in marine vertebrate genomics. Nature Reviews Genetics, 17, 523–534.

Lal MM, Southgate PC, Jerry DR, Zenger KR (2016) Fishing for divergence in a sea of connectivity: The utility of ddRADseq genotyping in a marine invertebrate, the black-lip pearl oyster Pinctada margaritifera. Marine genomics, 25, 57–68.

Lamichhaney S, Martinez Barrio A, Rafati N et al. (2012) Population-scale sequencing reveals genetic differentiation due to local adaptation in Atlantic herring. Proceedings of the National Academy of Sciences of the United States of America, 109, 19345–19350.

Laporte M, Pavey SA, Rougeux C et al. (2016) RAD-sequencing reveals within-generation polygenic selection in response to anthropogenic organic and metal contamination in North Atlantic Eels. Molecular Ecology, 25, 219–237.

Larson WA, McKinney GJ, Limborg MT et al. (2016) Identification of Multiple QTL Hotspots in Sockeye Salmon (Oncorhynchus nerka) Using Genotyping-by-Sequencing and a Dense Linkage Map. Journal of Heredity, 107, 122–133.

Larson WA, Seeb LW, Everett MV et al. (2014) Genotyping by sequencing resolves shallow population structure to inform conservation of Chinook salmon (Oncorhynchus tshawytscha). Evolutionary Applications, 7, 355–369.

Le Moan A, Gagnaire PA, Bonhomme F (2016) Parallel genetic divergence among coastal–marine ecotype pairs of European anchovy explained by differential introgression after secondary contact. Molecular Ecology, 25, 3187–3202.

Legrand JJ, Legrand Hamelin E, Juchault P (1987) Sex determination in Crustacea. Biological Reviews, 62, 439–470.

Lien S, Ben F Koop, Sandve SR et al. (2016) The Atlantic salmon genome provides insights into rediploidization. Nature, 533, 200–205.

Lischer HEL, Excoffier L (2012) PGDSpider: an automated data conversion tool for connecting population genetics and genomics programs. Bioinformatics, 28, 298–299.

Losada A (2014) Cohesin in cancer: chromosome segregation and beyond. Nature Reviews Cancer, 14, 389–393.

Lv J, Feng L, Bao Z et al. (2013) Molecular Characterization of RXR (Retinoid X Receptor) Gene Isoforms from the Bivalve Species Chlamys farreri. PloS one, 8, 096065.

Martin M (2011) Cutadapt removes adapter sequences from high-throughput sequencing reads. EMBnet journal, 17, pp. 10–12.

Martínez P, Viñas AM, Sánchez Let al. (2014) Genetic architecture of sex determination in fish: applications to sex ratio control in aquaculture. Frontiers in Genetics, 5, 152.

Mascher M, Wu S, Amand PS, Stein N, Poland J (2013) Application of Genotyping-by-Sequencing on Semiconductor Sequencing Platforms: A Comparison of Genetic and Reference-Based Marker Ordering in Barley H Candela, Ed,). PloS one, 8, 096065.

Mei J, Gui J-F (2015) Genetic basis and biotechnological manipulation of sexual dimorphism and sex determination in fish. Science China Life Sciences, 58, 124–136.

Miller AD, Rooyen A, Rašić G et al. (2016) Contrasting patterns of population connectivity between regions in a commercially important mollusc Haliotis rubra: integrating population genetics, genomics and marine LiDAR data. Molecular Ecology, 25, 3845–3864.

Moore JS, Bourret V, Dionne M et al. (2014) Conservation genomics of anadromous Atlantic salmon across its North American range: outlier loci identify the same patterns of population structure as neutral loci. Molecular Ecology, 23, 5680–5697.

Moore JS, N Harris Les, Kessel ST et al. (2016) Preference for nearshore and estuarine habitats in anadromous Arctic char (Salvelinus alpinus) from the Canadian high Arctic (Victoria Island, Nunavut) revealed by acoustic telemetry. Canadian Journal of Fisheries and Aquatic Sciences, 73, 1434–1445.

Mossman CA, Waser PM (1999) Genetic detection of sex-biased dispersal. Molecular Ecology, 8, 1063–1067.

Moura AE, van Rensburg CJ, Pilot M et al. (2014) Killer Whale Nuclear Genome and mtDNA Reveal Widespread Population Bottleneck During the Last Glacial Maximum. Molecular Biology and Evolution, 31, msu058–1131.

Narum SR, Buerkle CA, Davey JW, Miller MR, Hohenlohe PA (2013) Genotyping~by~sequencing in ecological and conservation genomics. Molecular Ecology, 22, 2841–2847.

Nayfa MG, Zenger KR (2016) Unravelling the effects of gene flow and selection in highly connected populations of the silver-lip pearl oyster (Pinctada maxima). Marine genomics, 28, 99–106.

Nikčević G, Savić T, Kovačević Grujičić N, Stevanović M (2008) Up~regulation of the SOX3 gene expression by retinoic acid: characterization of the novel promoter~response element and the retinoid receptors involved. Journal of Neurochemistry, 107, 1206–1215.

Ogden R, Gharbi K, Mugue N et al. (2013) Sturgeon conservation genomics: SNP discovery and validation using RAD-sequencing. Molecular Ecology, 22, 3112–3123.

Palti Y, Vallejo RL, Gao G et al. (2015) Detection and Validation of QTL Affecting Bacterial Cold Water Disease Resistance in Rainbow Trout Using Restriction-Site Associated DNA Sequencing H Wang, Ed,). PloS one, 10, 096065.

Pan Q, Anderson J, Bertho S et al. (2016) Vertebrate sex-determining genes play musical chairs. Comptes rendus biologies, 339, 258–262.

Pavey SA, Gaudin J, Normandeau E et al. (2015) RAD-sequencing Highlights Polygenic Discrimination of Habitat Ecotypes in the Panmictic American Eel. Current Biology, 25, 1666–1671.

Pecoraro C, Babbucci M, Villamor A et al. (2016) Methodological assessment of 2bRAD genotyping technique for population structure inferences in yellowfin tuna (Thunnus albacares). Marine genomics, 25, 43–48.

Perreault-Payette A, Muir AM, Goetz F, Sirois P, Perrier C, Normandeau E, Bernatchez L. 2016. Investigating the extent of parallelism in morphological and genomic divergence among lake trout ecotypes in Lake Superior. Molecular Ecology.

Picq S, McMillan WO, Puebla O (2016) Population genomics of local adaptation versus speciation in coral reef fishes (Hypoplectrus spp, Serranidae). Ecology and Evolution, 6, 2109–2124.

Poćwierz-Kotus A, Kijewska A, Petereit C et al. (2015) Genetic differentiation of brackish water populations of cod Gadus morhua in the southern Baltic, inferred from genotyping using SNP-arrays. Marine genomics, 19, 17–22.

Prugnolle F, De Meeûs T (2002) Inferring sex-biased dispersal from population genetic tools: a review. Heredity, 88, 161–165.

Pujolar JM, Jacobsen MW, Als TD et al. (2014) Genome-wide single-generation signatures of local selection in the panmictic European eel. Molecular Ecology, 23, 2514–2528.

Pusey AE (1987) Sex-biased dispersal and inbreeding avoidance in birds and mammals. Trends in Ecology & Evolution, 2, 295–299.

Reitzel AM, Herrera S, Layden MJ, Martindale MQ, Shank TM (2013) Going where traditional markers have not gone before: utility of and promise for RAD-sequencing in marine invertebrate phylogeography and population genomics. Molecular Ecology, 22, 2953–2970.

Ribas L, Robledo D, Gómez-Tato A et al. (2015a) Comprehensive transcriptomic analysis of the process of gonadal sex differentiation in the turbot (Scophthalmus maximus). Molecular and Cellular Endocrinology, 1–18.

Ribas L, Robledo D, Gómez-Tato A et al. (2015b) Comprehensive transcriptomic analysis of the process of gonadal sex differentiation in the turbot (Scophthalmus maximus). Molecular and Cellular Endocrinology, 1–18.

Rodríguez-Ezpeleta N, Bradbury IR, Mendibil I et al. (2016) Population structure of Atlantic Mackerel inferred from RAD-seq derived SNP markers: effects of sequence clustering parameters and hierarchical SNP selection. Molecular Ecology Resources.

Rougemont Q, Gagnaire P-A, Perrier C et al. (2016) Inferring the demographic history underlying parallel genomic divergence among pairs of parasitic and nonparasitic lamprey ecotypes. Molecular Ecology.

Rondeau EB, Minkley DR, Leong JS et al. (2014). The genome and linkage map of the northern pike (Esox lucius): conserved synteny revealed between the salmonid sister group and the Neoteleostei. PLoS One, 9, 096065.

Sakamoto T, Danzmann RG, Gharbi K et al. (2000) A Microsatellite Linkage Map of Rainbow Trout (Oncorhynchus mykiss) Characterized by Large Sex-Specific Differences in Recombination Rates. Genetics, 155, 1331–1345.

Sivasubramaniam S, Sun X, Pan Y-R, Wang S, Lee EYHP (2008) Cep164 is a mediator protein required for the maintenance of genomic stability through modulation of MDC1, RPA, and CHK1. Genes & Development, 22, 587–600.

Sodeland M, Jorde PE, Lien S et al. (2016) “Islands of Divergence” in the Atlantic Cod Genome Represent Polymorphic Chromosomal Rearrangements. Genome Biology and Evolution, 8, 1012–1022.

Stockwell BL, Larson WA, Waples RK et al. (2015) The application of genomics to inform conservation of a functionally important reef fish (Scarus niger) in the Philippines. Conservation Genetics, 1–11.

Sutherland B, Gosselin T, Normandeau E et al. (2016) Salmonid chromosome evolution as revealed by a novel method for comparing RADseq linkage maps. Genome Biology and Evolution.

Team RC (2013) R: A language and environment for statistical computing.

Tepolt CK, Palumbi SR (2015) Transcriptome sequencing reveals both neutral and adaptive genome dynamics in a marine invader. Molecular Ecology, 24, 4145–4158.

Therkildsen NO, Hemmer-Hansen J, Als TD et al. (2013) Microevolution in time and space: SNP analysis of historical DNA reveals dynamic signatures of selection in Atlantic cod. Molecular Ecology, 22, 2424–2440.

Timusk ER, Ferguson MM, Moghadam HK et al. (2011) Genome evolution in the fish family salmonidae: generation of a brook charr genetic map and comparisons among charrs (Arctic charr and brook charr) with rainbow trout. BMC genetics, 12, 68.

Woram RA, Gharbi K, Sakamoto T et al. (2003) Comparative genome analysis of the primary sex-determining locus in salmonid fishes. Genome research, 13, 272–280.

Wright S (1931) Evolution in Mendelian populations. Annals of Eugenics, 16, 97–159.

Xu T, Sun J, Lv J et al. (2016) Genome-wide discovery of single nucleotide polymorphisms (SNPs) and single nucleotide variants (SNVs) in deep-sea mussels_ Potential use in population genomics and cross-species application. Deep-Sea Research Part II, 1–9.

Yano A, Nicol B, Jouanno E et al. (2013) The sexually dimorphic on the Y-chromosome gene (sdY) is a conserved male~specific Y~chromosome sequence in many salmonids. Evolutionary Applications, 6, 486–496.

Zhang B-D, Xue D-X, Wang J et al. (2016) Development and preliminary evaluation of a genomewide single nucleotide polymorphisms resource generated by RAD-seq for the small yellow croaker (Larimichthys polyactis). Molecular Ecology Resources, 16, 755–768.

